# Multiple random phosphorylations in clock proteins provide long delays and switches

**DOI:** 10.1101/2020.06.07.138438

**Authors:** Abhishek Upadhyay, Daniela Marzoll, Axel Diernfellner, Michael Brunner, Hanspeter Herzel

## Abstract

Theory predicts that self-sustained oscillations require robust delays and nonlinearities (ultrasensitivity). Delayed negative feedback loops with switch-like inhibition of transcription constitute the core of eukaryotic circadian clock. The kinetics of core clock proteins such as PER2 in mammals and FRQ in *Neurospora crassa* is governed by multiple phosphorylations. We investigate how multiple, slow and random phosphorylations control delay and molecular switches. We model phosphorylations of intrinsically disordered clock proteins (IDPs) using conceptual models of sequential and distributive phosphorylations. Our models help to understand the underlying mechanisms leading to delays and ultrasensitivity. The model shows temporal and steady state switches for the free kinase and the phosphoprotein. We show that random phosphorylations and sequestration mechanisms allow high Hill coefficients required for self-sustained oscillations.

## 1. Introduction

Life on earth in forms of cyanobacteria, algae, fungi, plants and animals has evolved 24 hour periodicities called circadian clocks. This helps them to anticipate rhythmic environmental cues such as light, temperature and nutrients [1–6]. Circadian clocks regulate a wide variety of molecular and physiological processes [7–10].

Circadian oscillators are based on a transcription-translation feedback loops (TTFLs). A delayed negative feedback loop is central to the gene regulatory network [11–13]. For example, the negative feedback loop of the fungal clock contains the negative element FREQUENCY [FRQ], which inhibits its own expression via inhibition of the circadian transcription factor White Collar Complex (WCC). FRQ is an intrinsically disordered protein (IDP) progressively hyperphosphorylated mainly by CK1a (Casein Kinase 1a). Hyperphosphorylation eventually leads to functional inactivation and degradation of FRQ allowing the WCC to reinitiate a new cycle [14]. This design principle is conserved in other frequently used model systems. Fruit flies and animal clocks are also made up of the inhibitors, kinases and transcriptional activators. In mammalian clocks, PERIOD proteins [PER1, PER2, PER3], and CRYPTOCHROME proteins [CRY1, CRY2]) inhibit their own expression. PER2 is also an IDP as FRQ and is phosphorylated by CK1 [15–17].

Recent experiments suggest that many sites on both FRQ (about 100 sites) and PER2 (about 60 sites) are phosphorylated over the course of many hours at seemingly random manner [18–21]. Moreover their respective activators also get phosphorylated [15,22,23]. Phosphorylations govern nuclear translocation [24], complex formations [25,26], inactivation of transcription and stability [27,28].

Oscillator theory predicts that self-sustained circadian clocks require long delays and nonlinearities such as switches [29,30]. This raises two theoretical questions: (1) How can multiple random phosphorylations produce long delays? (2) What are the underlying switch mechanisms?

According to mathematical theory a delay of negative feedback loops governs the period [29,31]. Under quite general assumptions the delay is in the range between a 1/4 and 1/2 of the oscillator period [32,33]. If gene expression and inhibitor formation last about an hour, periods of 2 to 4 hours can be expected. Indeed, several TTFLs exhibit periods of a few hours including somite formation [34], NF-kB rhythms [35,36] and p53 pulses [37]. Circadian rhythms have much longer periods of about 24 hours. This implies that the associate delays last at least 6 hours. Several processes like transcription, translation, nuclear transport, post-translational modifications, mRNA decay, and proteasomal degradation may contribute to the needed delay in circadian rhythms [15,24]. It has been suggested that also the multiple phosphorylations contribute significantly to the required delay [38,39].

In order to generate self-sustained rhythms (“limit cycles”) nonlinearities are necessary in addition to delays [40,41]. In many models switch-like inhibitions are postulated [42,43]. Here we explore how multiple random phosphorylation contribute to the generation of switch-like behaviour.

Fig 1 depicts the assumptions underlying our modelling approach: There is clear experimental evidence for a slow and random phosphorylation of the clock phosphoproteins [18,44]. Fig 1A depicts gradual phosphorylation in *Neurospora*. FRQ is stabilized by the FRQ-INTERACTING HELICASE (FRH) and forms the FFC complex with the kinase CK1a [14,45,46]. FRQ undergoes slow, seemingly random multiple phosphorylations predominantly by CK1a [18,47–49]. At an overcritical phosphorylation level the complex gets inactivated in a switch-like manner [23,27,50,51].

**Figure 1.**
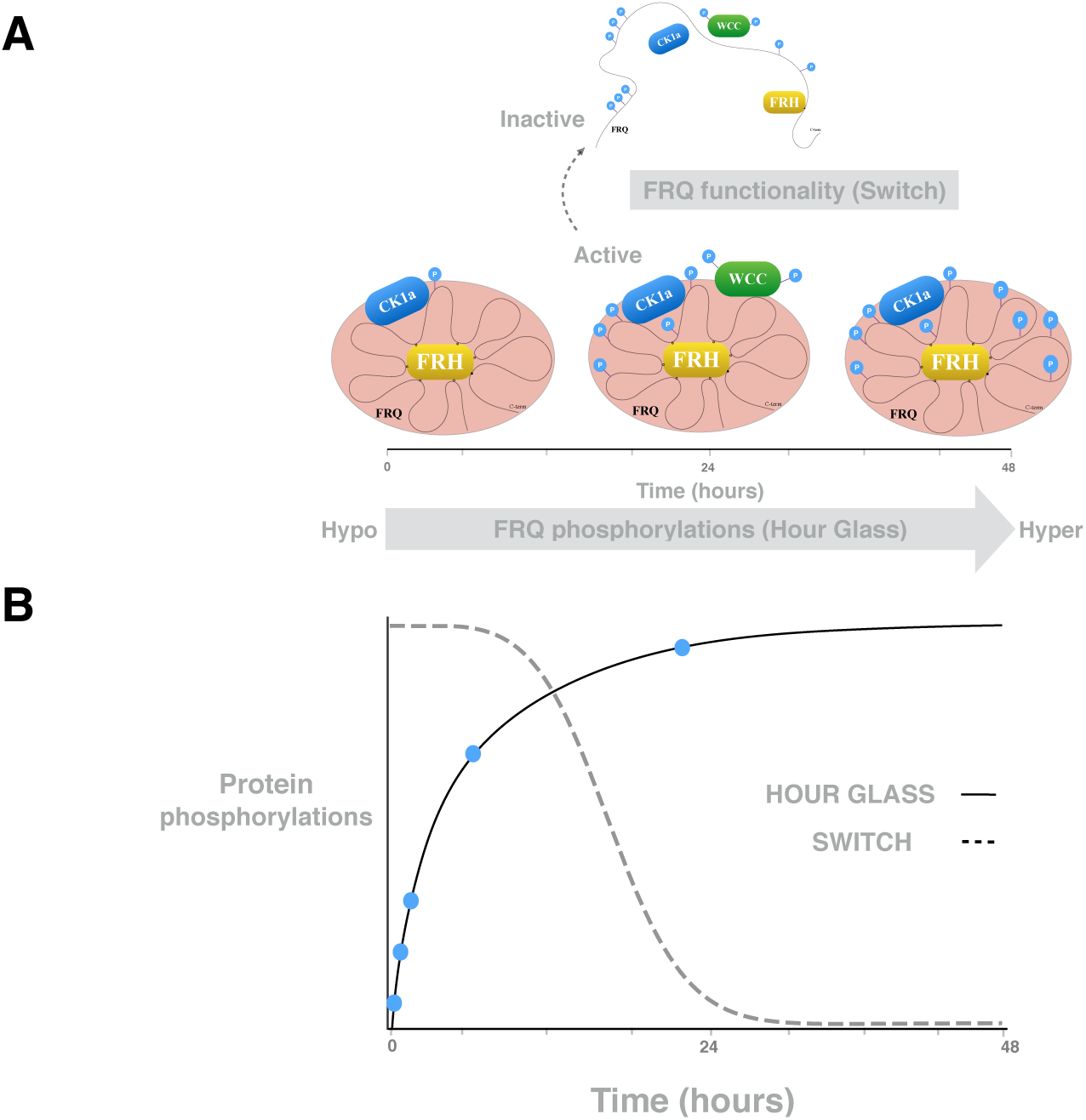
Phosphorylation hourglass and activity switch phosphoproteins: **(A)** As an example, the FRQ protein undergoes multiple phosphorylations. A phosphoswitch governs the activity status of the complex.**(B)** The solid line represents the simulated phosphorylations in Fig 7 and suggests a slow and saturated phosphorylation. The dashed line represents the simulated switch induced by hyperphosphorylated protein (compare Fig 7).

Since detailed kinetic data of these processes are missing we explore in generic models how multiple random phosphorylations can reproduce long robust delays and switch-like behaviour. Fig 1B illustrates our conceptional modelling approach: We simulate slow saturated phosphorylation (solid line) leading to a switch-like inactivation at critical phosphorylation levels.

## 2. Results

### 2.1. Linear processive phosphorylations provides delays

As discussed above, PER2 and FRQ are core clock phosphoproteins with up to 100 phosphorylation sites [28,52]. Recent *in vitro* and *in vivo* experiments show that in *Neurospora* about 100 FRQ sites are phosphorylated over more than a circadian day (up to 48 hours) in a seemingly random manner [18]. However the detailed functions of increasing phosphorylation levels in circadian timekeeping are not well understood.

Inspired by these observations, we introduce conceptual models of multiple phosphorylations. We denote the simulated clock phosphoprotein by “F” and the associated kinase by “C”. For simplicity, we start with just 4 phosphorylation sites. In A1 we show the corresponding reaction scheme and the associated linear differential equations describing protein turnover and processive phosphorylation. We do not explicitly consider phosphatases throughout. Phosphatases are found to be permanently available in cells and it is basically the ratio of kinase and phosphatase which takes the reaction forward [53,54]. Fig 2 shows the increasing levels of different phosphorylations in the linear model. Note that we introduce just two parameters: a production and phosphorylation rate k and a degradation rate kd. We have chosen parameter values that reproduce the measured full phosphorylation after about 48 hours. [18,44]. Thus we can focus on delays, amplitudes and waveforms quantified by fits of Hill-functions (see section 4.2 Materials and methods)

**Figure 2.**
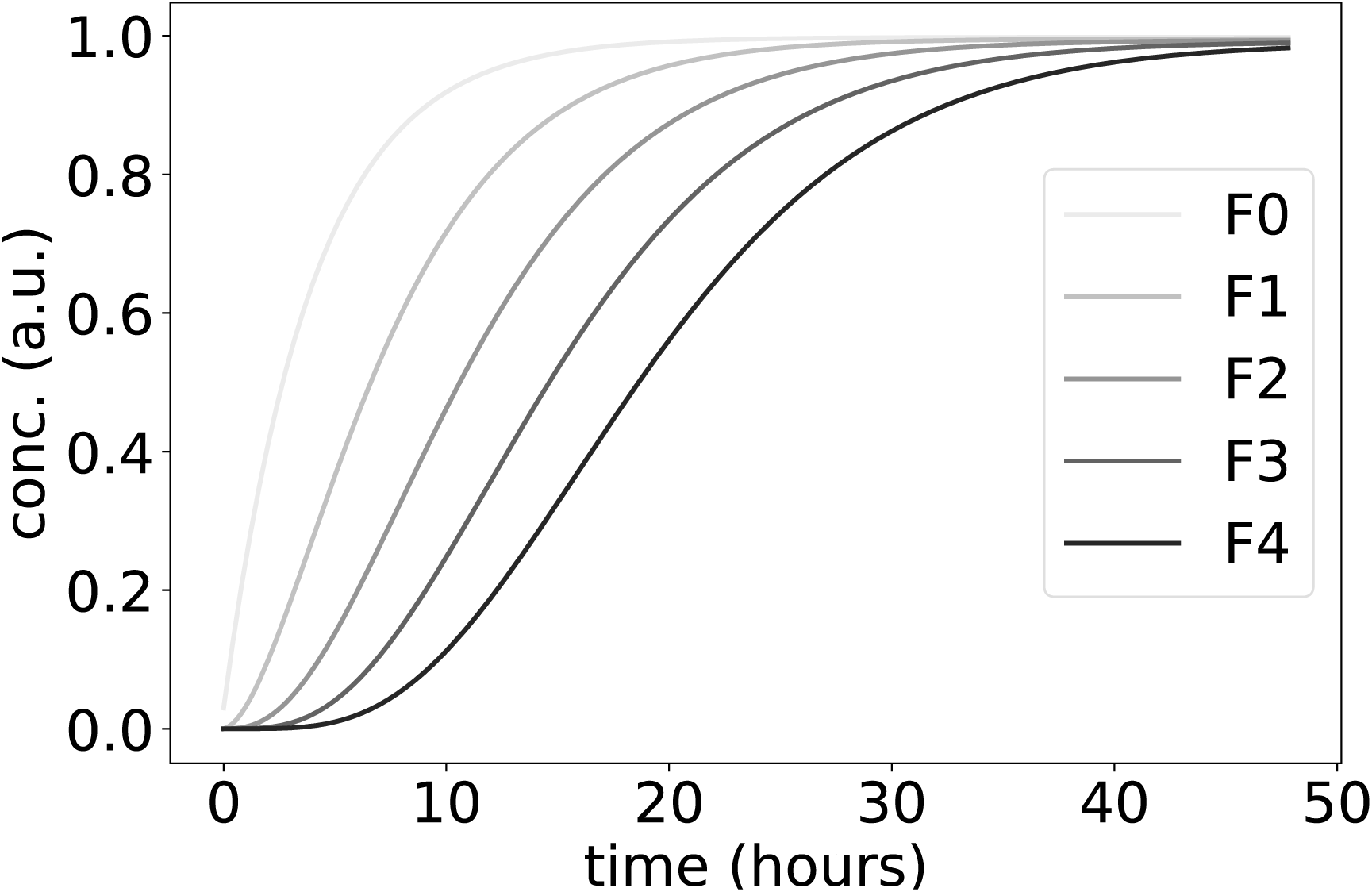
Modeling a delayed switch based on multiple phosphorylations: The simulations start with unphosphorylated protein F. The normalized total number of Fk with k=1,..,4 phosphorylations increases with time delays and hyperphosphorylated species (such as n = 4) accumulate in an ultrasensitive, switch-like manner.

The curves for phosphoprotein F exhibited delays and an accumulation of fully phosphorylated F4 in a switch-like manner (Fig 2). Therefore even linear phosphorylations alone can provide delays. A delayed switch based on multiple phosphorylations can serve as the basic hourglass mechanism hypothesized in Fig 1. Note, that these steps required no explicit nonlinearity in the model.

### 2.2. Nonlinear models of distributive phosphorylations enhance ultrasensitivity

Kinases bind to substrates and could phosphorylate sites while staying bound (processive mechanism). Alternatively, the kinase may bind and unbind, so that next phosphorylation first requires rebinding of a kinase molecule (distributive mechanism) [53]. Distributive enzyme kinetics may lead to ultrasensitive responses in protein phosphorylations [55,56]. Note that in these studies ultrasensitivity is quantified using input-output relations. Typically steady states of phosphorylation levels are studied as a function of ligand concentrations or kinase levels. Motivated by our temporal switch in Fig 1, we focus on the ultrasensitive increase of phosphorylation with time. In this section we include the formation and dissociation of FC complexes in order to study the role of enzyme sequestration.

Our models are motivated by experimental observations regarding multiple slow and partly random phosphorylations of clock proteins [18,44]. We emphasize, however, that the following model versions do not reflect the full complexity of clock protein interactions as phosphorylations can affect stability and complex formation in a complicated manner [25,27]. If the kinase stays bound to the substrate it is termed processive phosphorylation (see A1). Dissociation of C and rebinding to other F molecules could be represented as distributive mechanism.

The simulations below are somewhat simplistic and describe generic substrate-enzyme dynamics. For example, we start simulations with low levels of F and of recruited enzyme C. Kinase C can be recruited to unphosphorylated and partially phosphorylated species of protein F (F0, F1, F2 and F3). The bound kinase can then be sequestered (F1C, F2C,..) which slows down the progressive phosphorylation kinetics. Eventually F is further phosphorylated and C dissociates, yielding the next phospho-species.

A2 shows the turnover of F (Fk) and non-sequestered and sequestered CF complexes, Cf and FkC, respectively. Using mass-action kinetics, this scheme is directly translated into a system of nonlinear ordinary differential equations (ODEs) [57–59]. The equations describe 10 time-varying concentrations of (phosphorylated) F, complexes with the kinase and turnover of C. The 5 kinetic parameters in the ODEs are enumerated as Pp and Pc, k1, k2 and kd (see A2) respectively for production of protein and kinase, rates of phosphorylation and degradation.

Fig 3 shows the production, dissociation and phosphorylation of F and the formation of FC complexes starting at zero levels. F0 denotes the unphosphorylated F protein whereas F1, F2, F3, F4 represent increasing phosphorylation levels. Ct and Cf denote the amounts of total and free kinase, respectively. The phosphorylated F species accumulate in a sigmoidal manner with the expected delays. There is an initial sharp peak of FC complexes and of free enzyme Cf but later most of the enzyme molecules are “sequestered”. A5 shows the Hill coefficients for phosphorylated Fk (F0…F4) calculated along the lines of [60,61]. In comparison to linear phosphorylations shown in Fig 2, nonlinear phosphorylations provide higher Hill coefficients. Note, however, that the amplitudes decrease drastically for higher phosphorylation levels.

**Figure 3.**
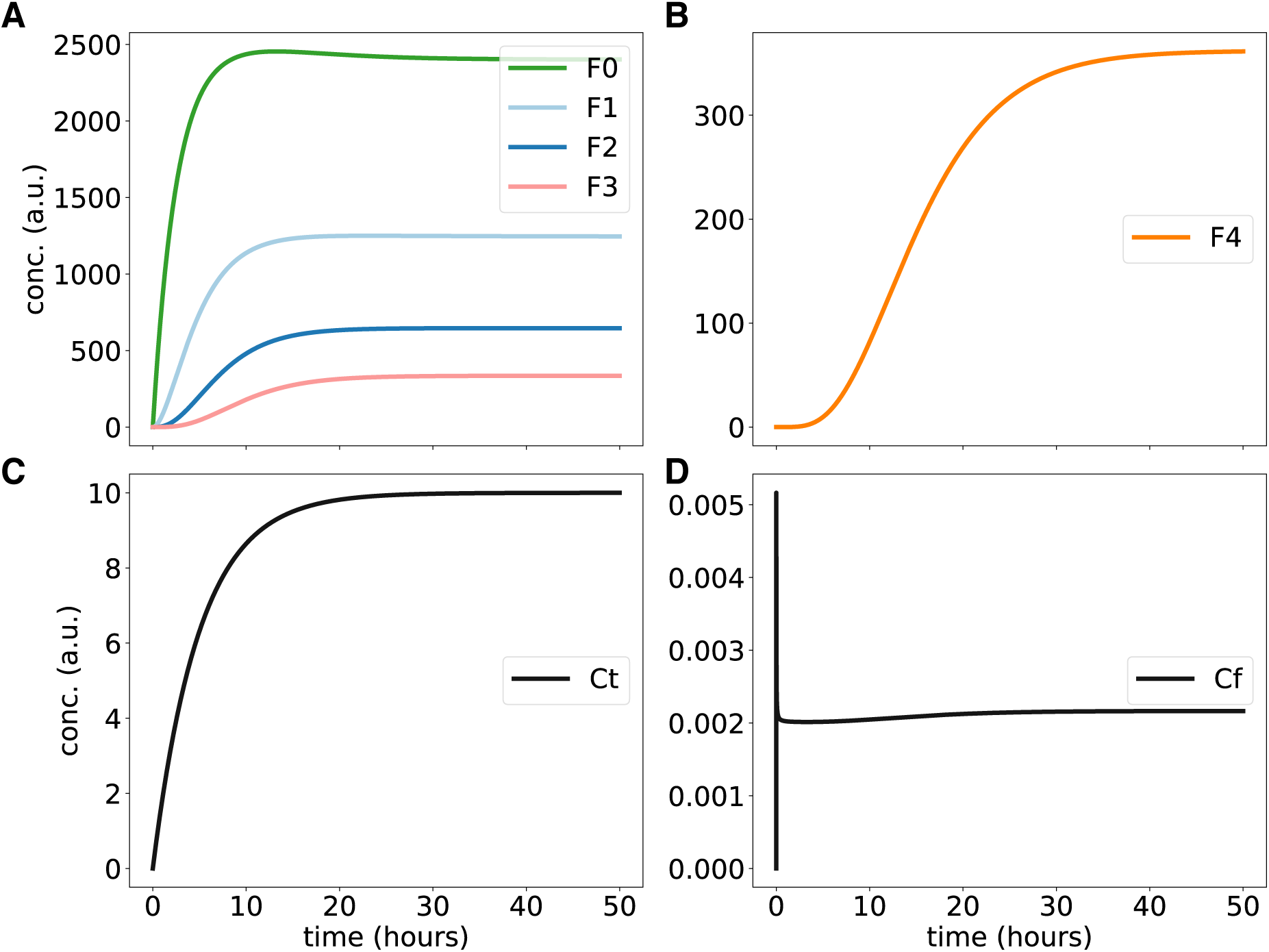
Kinetics of nonlinear distributive phosphorylations: (A) and (B) show steep curves of hypo- and hyper-phosphorylated F proteins over time. Graph (C) and (D) display total and free kinase concentrations, respectively.

### 2.3. Sequestration can generate overshoots and threshold behaviour

It was shown in the preceding section that sequestration can enhance switch-like behaviour. In Fig 3, the simulated amount of free enzyme was relatively small compared to the phosphoprotein F and thus most of the enzyme was sequestered. It is known, however, that kinase levels are quite high in *Neurospora* [62]. Consequently, we study here the effects of enlarged kinase levels. It turns out that higher kinase levels lead to overshoots and a sharp threshold.

In Fig 4 the levels of phosphorylated proteins F1, F2, and F3 increase initially as in Fig 3. After about 10 hours, however, they decay to quite small values. In parallel, the fully phosphorylated protein F4 reaches high levels (see Fig 4B). If full phosphorylation reaches saturation the amount of free kinase is increasing suddenly. Magnifications reveal that the apparent “kinks” in the time-courses are smooth curves (see A6).

**Figure 4.**
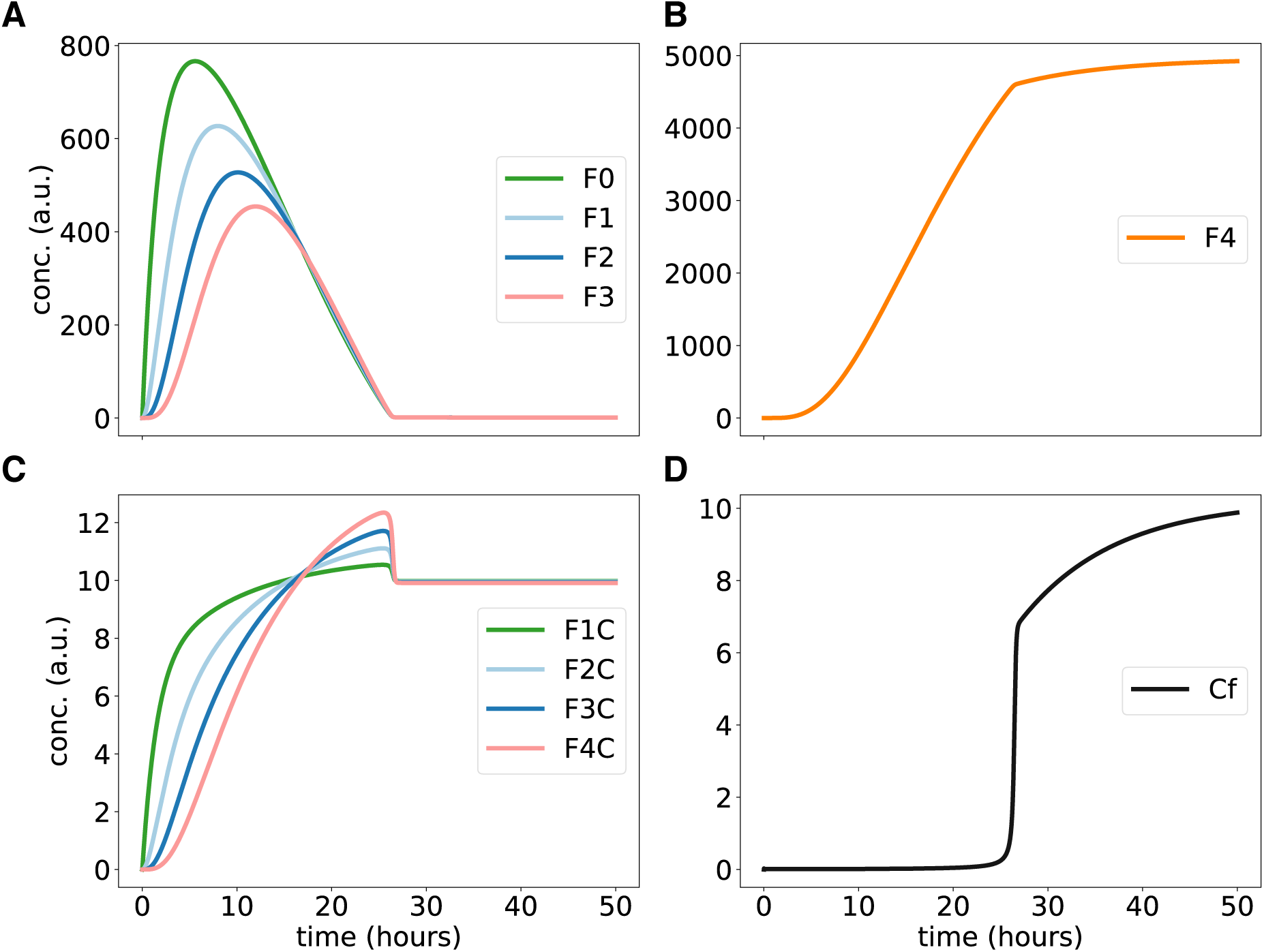
Simulating nonlinear phosphorylations at higher enzyme levels: (A) and (B) show time-courses of phosphorylated F proteins. Graph (C) shows the concentrations of FC complexes over time. (D) shows the temporal switch of free kinase Cf after release from sequestration. The dynamics of total kinase Ct, not shown here, is similar as in Fig 3.

Note, that Figs 3 and 4 refer both to our model described in A2. The drastic differences are simply due to 5-fold increase of enzyme production. The overshoot in Fig 4 reflects the fast initial production of F0 and phosphorylation of F1, F2 and F3. Later an equilibrium is reached with lower levels of intermediate phosphorylations.

In summary, for higher kinase levels threshold behaviour arises reflecting the sequestration of enzymes by different species of phosphorylated proteins. Note, that the initial sharp increase of free protein levels is based on our somewhat artificial initial condition of zero enzyme levels. In vivo the equilibrium between free and bound proteins is reached more quickly due to the omnipresence of kinases.

### 2.4. Steady state switches due to increasing enzyme levels

So far, we have characterized in Figs 3 and 4 two specific values of enzyme production Pc (marked by arrows in Fig 5). Fig 5 illustrates that a systematic variation of enzyme levels induces nonlinear dependencies including transitions between different regimes. For low values of Pc we get power-law increases of free kinase (exponent =2.85) and of sequestered protein (exponent = 4.29). Note, that the model contains just bilinear terms. Thus, sequential phosphorylation and sequestration leads to higher exponents as described earlier in other systems [56,60,63]. As discussed above the generation of such input-output switches was studied extensively over the last decades [64–66]. Even though our model can reproduce such nonlinearities, our focus are temporal switches (compare Figures 1 to 4) relevant for circadian rhythm generation.

**Figure 5.**
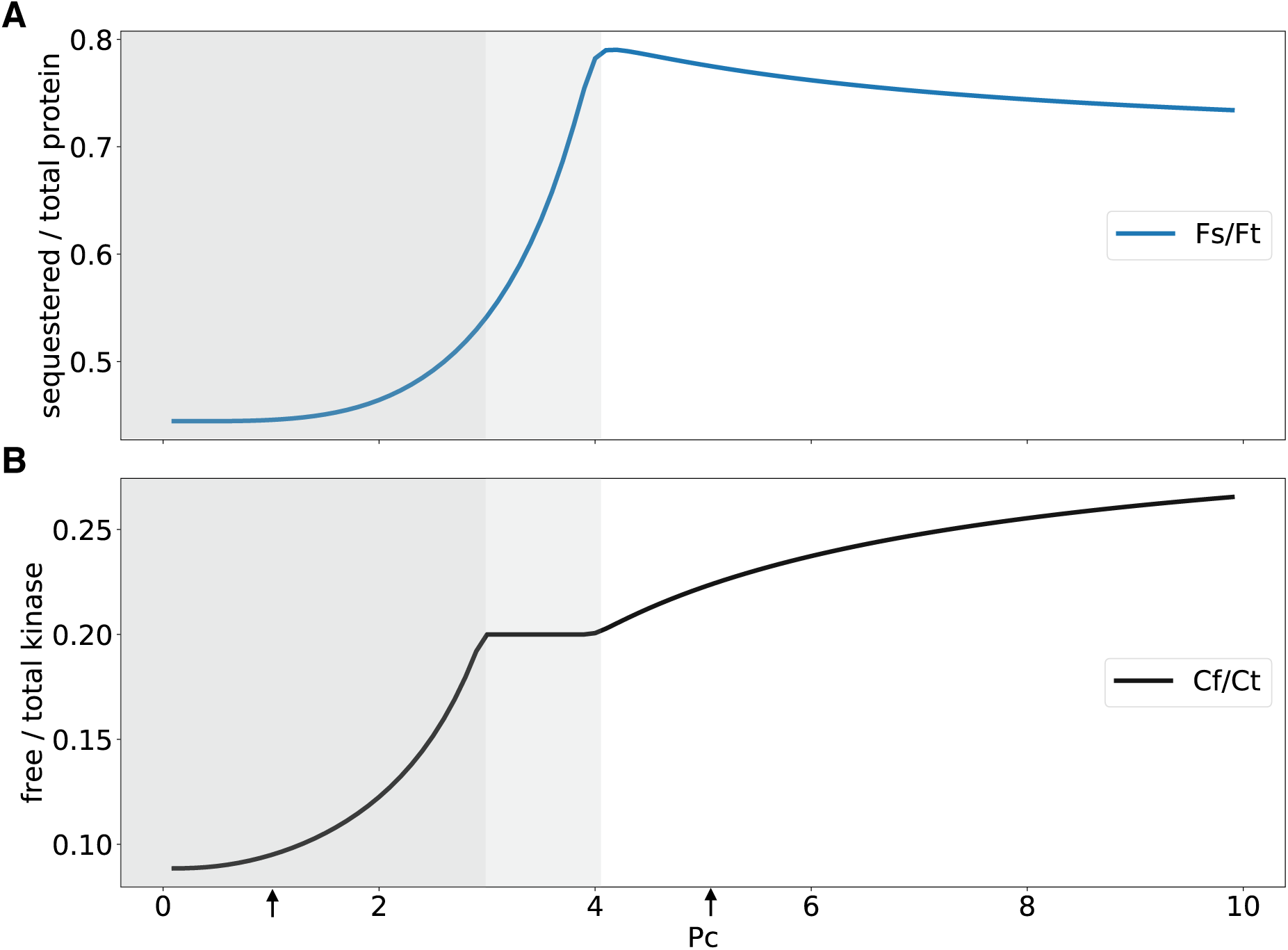
Simulating sequestration based transitions: (A) shows the ratio of sequestered to total F for varying kinase production. (B) shows the ratio of free to total kinase for increasing kinase production.

Inspecting Fig 5, we find that at Pc = 3 sequestration by increasing protein F levels flattens the curve of free kinase due to sequestration. Beyond Pc = 4 (white part of the graph) the amount of sequestered protein starts to decline and the release of enzyme leads to more free kinase *C*_*f*_.

In summary Fig 5 shows that our nonlinear model generates, in addition to delayed temporal switches and threshold behaviour, interesting nonlinearities of steady states governed by sequestration.

### 2.5. Random phosphorylations provide large amplitudes of intermediate phosphorylations

In signaling cascades many phosphorylation sites carry specific functions for activations or complex formations. Up to 100 phosphorylations of intrinsically disordered proteins (IDPs) such as FRQ or PER2 might control cellular processes differently [15]. FRQ proteins have a positively charged N-terminal part and a negatively charged C-terminal part. Initial phosphorylations appear to stabilize a closed conformation whereas progressive hyperphosphorylations favours an open conformation potentially via charge repulsion [51]. Thus, the overall number of phosphorylated sites can govern stability, complex formation or nuclear translocation [18,49].

There is some evidence that FRQ is phosphorylated by CK1a in a seemingly random opportunistic manner [18,52]. Hypothetically, the highly flexible FRQ protein can form almost random contacts to the active site of CK1a allowing phosphorylation [62]. Consequently we simulate now instead of processive phosphorylations (compare A1) random phosphorylations as shown in A3. Note, that for up to 100 phosphorylations the number of differently phosphorylated molecules is astronomically large and exceeds the number of FRQ molecules in a cell and even the number of bacteria on earth [67].

In order to keep simulations with differential equations feasible we introduce new variables F1, F2, etc. These variables lump all F molecules with 1, 2, … phosphorylations. In other words, Fk represents the 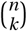 molecules with k out of n phosphorylations. These lumped variables seem reasonable since complex formation of clock proteins are influenced by the number of phosphorylations [18,25]. For example, an overcritical number of phosphorylations destabilizes the FFC complex and makes FRQ accessible to the E3 ligase FWD1 (F-box/WD-40 repeat-containing protein-1) [18,68,69].

Our resulting nonlinear random model (see A4) contains altogether just 2n+2 differential equation making even simulations with n=100 phosphorylations feasible. For a direct comparison with the previous models we show in Fig 6 simulations for n=4. It turns out that the delayed switch with Hill coefficient around 3 are found in these simulations as well (compare A5). Fig 6 displays a new feature of random models - the amplitudes of intermediate states do not decay monotonously. For instance, F1 and F2 have fairly high levels. This property of random models can be traced back to prefactors in the lumped equations. In other words, the combinatorial explosion of molecule types with intermediate phosphorylation numbers enhances the growth of certain Fk levels.

**Figure 6.**
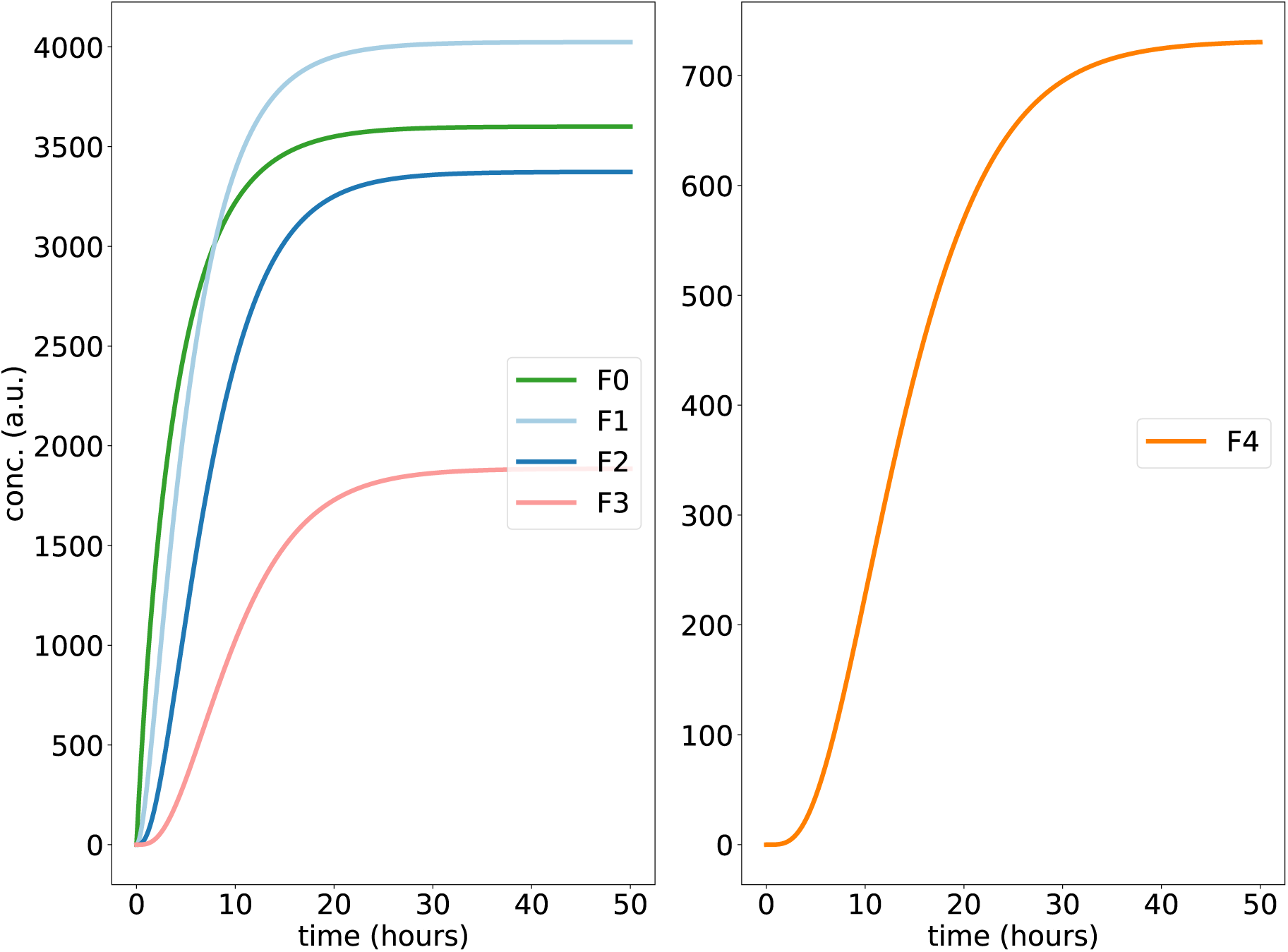
Simulating nonlinear random phosphorylations: Graphs show time-courses of hypo, medium, hyperphosphorylated and fully phosphorylated F proteins for n=4 phosphorylations.

Fig 7 shows representative time-courses of a random model with up to 100 phosphorylations. It turns out that we find delayed temporal switches with high amplitudes in particular at intermediate phosphorylation levels.

**Figure 7.**
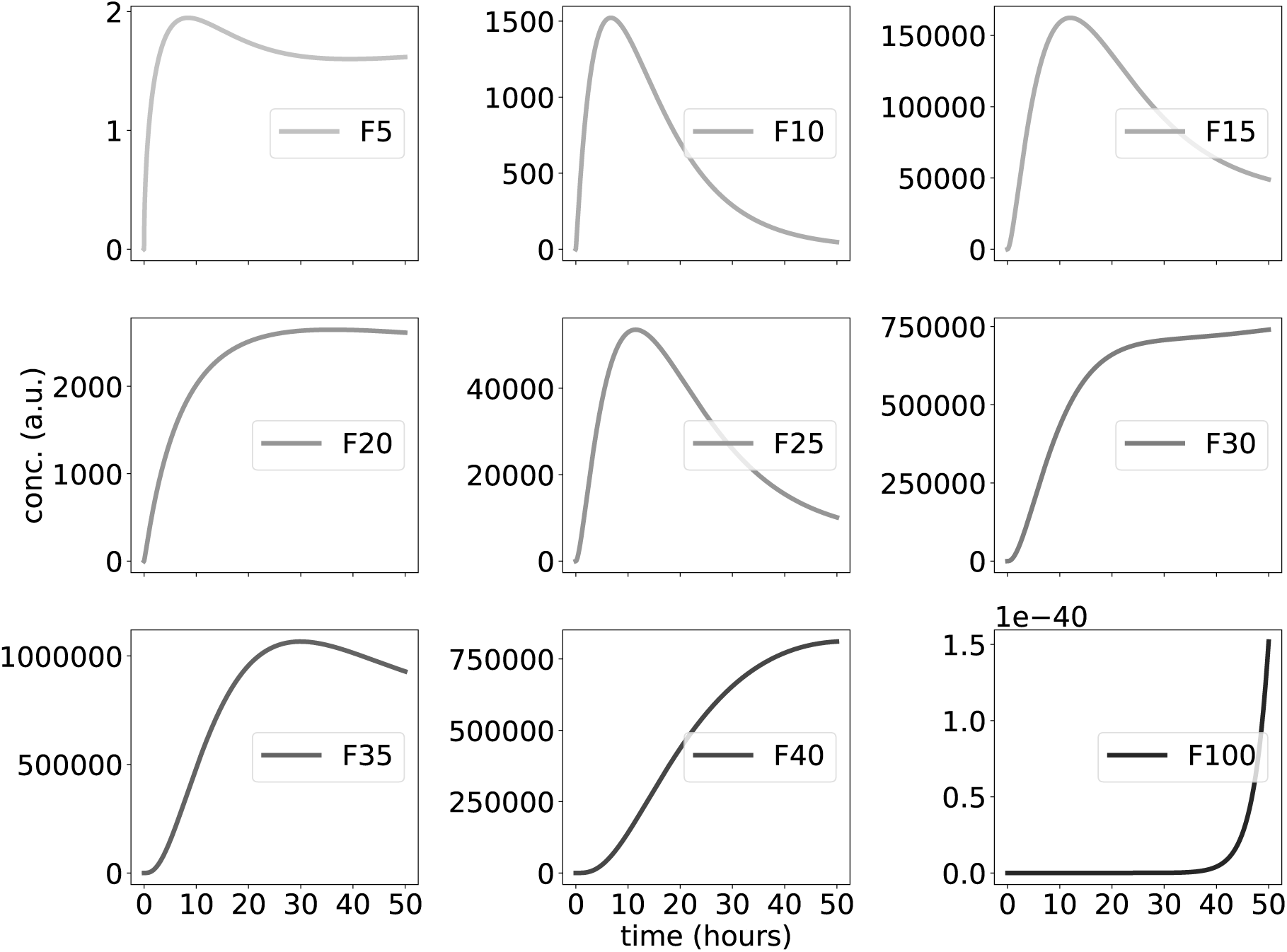
Large-scale simulations of nonlinear, random phosphorylations: Graphs show time-courses of hypo, medium, hyperphosphorylated and fully phosphorylated F proteins for up to n=100 phosphorylation sites. Note, that the dashed line in Fig 1 (inactivation of F due to phosphorylations) is adapted from 1 – F40 from this figure.

Note, that the widely varying amplitudes can not be directly related to experimental data. Nevertheless, there are generic features of the model that do not depend upon sensitivity to chosen parameter values (compare also Fig 8): hypophosphorylated proteins increase quickly up to medium levels. Levels of medium phosphorylations (F30…F40) exhibit ultrasensitive increases to fairly large amplitudes. Fully phosphorylated proteins are found only at tiny levels.

**Figure 8.**
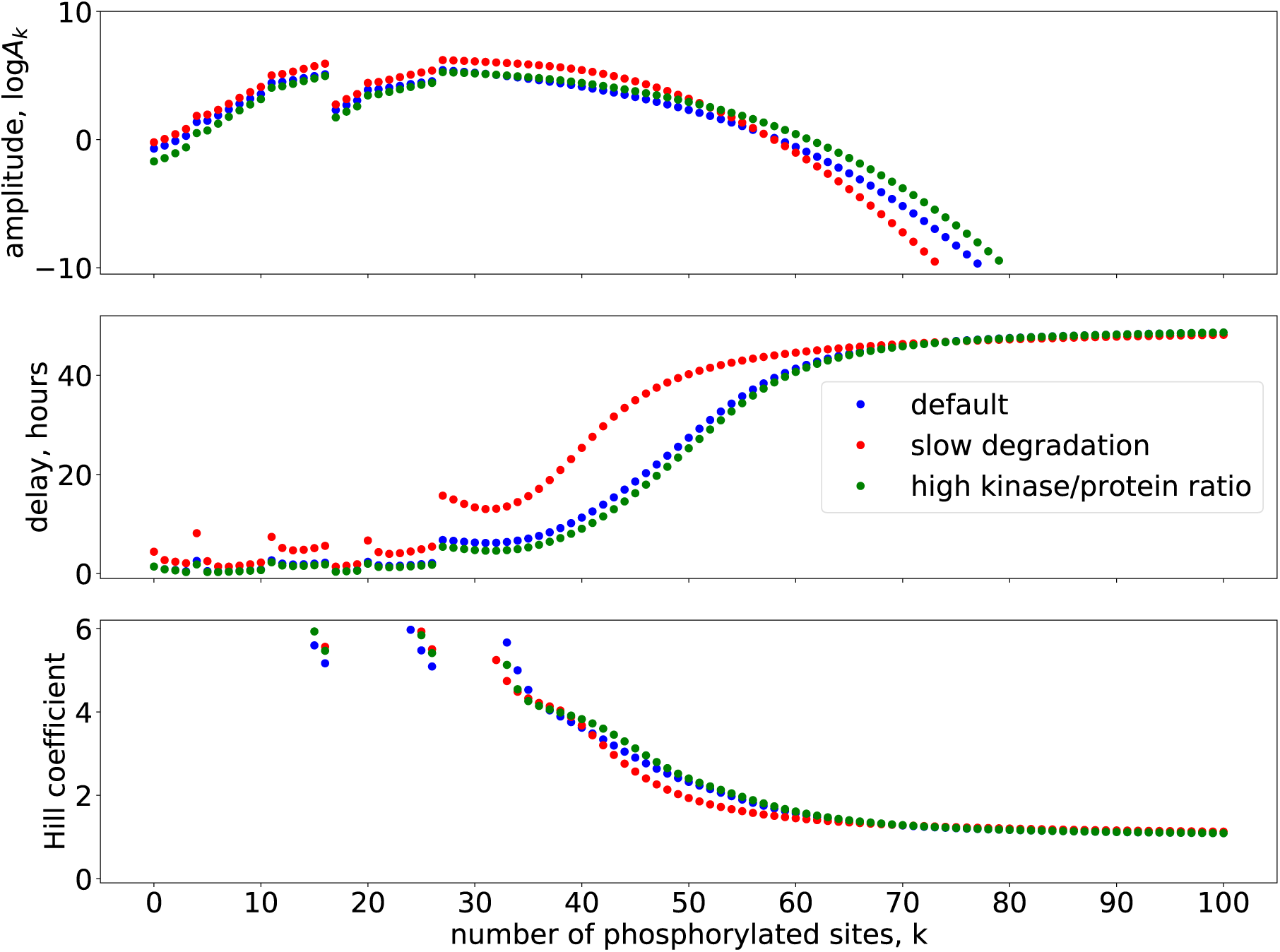
Intermediate phosphorylation levels exhibit large amplitudes, delays and ultrasensitivity: (A) shows log10 values of amplitude for phosphorylated species. (B) depicts delays and (C) shows Hill coefficients for phosphorylated species.

In Fig 8A we use a logarithmic scale to cover these drastic amplitude differences. The graph reveals that for different parameters intermediate levels of phosphorylations exhibit quite large amplitudes. The steps reflect changes in waveforms (see Fig 7) due to sequestration effects.

In addition to large amplitudes mediators of cellular switches require robust delays and ultrasensitivity. The graphs in Fig 8B and 8C show that intermediate phosphorylation levels obey these properties for different parameters constellations.

In order to test the robustness of these features we analyzed 10-fold reductions of the degradation rate (red) and a 10-fold increase of the enzyme/substrate ratio (green). It turns out that the main properties - huge amplitudes at intermediate phosphorylation levels, long delays, and Hill coefficients above 3 - are robust with respect to large parameter variations.

## 3. Discussion

Self-sustained circadian oscillations require long delays and nonlinearities (“switches”) [29,30]. Recent experiments suggest that slow and seemingly random phosphorylations of intrinsically disordered clock proteins control stability and function of clock protein complexes such as FFC in *Neurospora* and PER:CRY in mammals [18,25,70]). Since only few detailed quantitative data are available we compared several generic models describing linear processive phosphorylation, nonlinear distributive phosphorylation, and random phosphorylation. We find that long delays are robustly achieved by long reaction chain lengths and slow degradation (Figs 2 to 4). Sequestration enhances the formation of temporal switches with high Hill coefficients (A5). Thus our simulations support our hypothesis sketched in Fig 1 that multiple random phosphorylation can provide robust delayed switches.

Our theoretical approach is based on a few experimental observations that help to constrain our generic models. Interestingly, most of the data suggest quite similar principles in the *Neurospora* clock and mammalian clocks. In both cases intrinsically disordered proteins (FRQ and PER, respectively) with multiple random phosphorylations are central players within the negative transcriptional-translational feedback loop [21,62,71,72]. Below we focus on the *Neurospora* clock.

The half-life of the FRQ protein is about 3-5 hours [73]. About 100 phosphorylation sites have been identified using isotope labelling and mass spectrometry. The priming-independent phosphorylation of non-consensus sites on FRQ by CK1a seems to be slower than 5 sites per hour [54]. Priming by other kinases is relatively fast and appears to be less essential for the principle function of the core clock. However, other kinases might be relevant for entrainment and temperature compensation not discussed in this paper [18].

Fig 1 illustrates the important role of multiple phosphorylations. FRQ has a positively net charged N-terminal part, a negatively net charged C-terminal part, and a central part involved in proteasomal degradation by FWD1 [51,68,73]. Initial phosphorylation of the C-terminal part early in the circadian day has a stabilizing effect [18,74]. Subsequent phosphorylation of the N-terminal part destabilized the FFC complex allowing degradation. Thus, if charge repulsion were to govern stability and function, the number of phosphorylations would play a central role.

These experimental findings are reviewed in recent publications [18,23,74] and provide the framework of our models. We do not fit individual parameters to the sparse quantitative data but we adapt the models design to the observations listed above. For example, the number of phosphorylations, the degradation rates, and the central role of casein kinase are consistent with the data. Since quantitative details of binding and unbinding of CK1a to FRQ are not known, we simulated two mechanisms: linear processive phosphorylations (A1 and Fig 2) and distributive phosphorylations (A2, A3, Figures 3 to 8). At a first glance, the distributive mechanism seems not consistent with Fig 1 showing a relatively stable complex involving FRQ and CK1a. Indeed, CK1a is first recruited to the FRQ-CK1a-domains (FCD1 and 2) of FRQ [74]. Then the active site of bound CK1a can phosphorylate step by step the FRQ protein. This could happen at the same FRQ molecule (processive) or after dissociation and rebinding at another FRQ molecule [51]. Such reoccurring binding events could be simulated by our approach using the distributive mechanism. We start our simulations with zero levels of protein and enzyme. This allows to follow the kinetics of complex formations, sequestration, and phosphorylations.

Our conceptual models could be extended in future studies by more detailed features of protein dynamics [18]. For example, the number of phosphorylations influences stability and dissociation constants implying dependencies of model parameters on the number of phosphorylations k [74]. In our A7 we show that decreasing phosphorylation rates and varying stability do not change our main results. In future studies, stabilizing effects of FRH [46], interactions with WCC [23], and degradation assisted by FWD1 [69] could be incorporated.

We emphasize, that our focus on multiple random phosphorylations neglects other essentials of the transcriptional-translational feedback loop (TTFL) modeled in detail elsewhere [26,75–78]. Nevertheless, delayed switch-like behavior due to slow random multiple phosphorylation seems to be central element in circadian rhythm generation. A delayed switch due to multiple phosphorylation is a robust design principle that could be relevant also in other biological systems such as ligand specificity, nuclear import, DNA binding in T-cells [53,79,80], timing of critical transitions in cell cycle [81], regulation of sleep-wake homeostasis in mice [82], Familial advanced sleep-phase syndrome (FASPS) in humans [24], phototactic sensitivity in green algae [83], and reproductive fitness in cyanobacteria [84].

## 4. Materials and methods

### 4.1. Numerics

All the simulations have been performed on a Spyder Python 3.4 platform. Simulations resulting in Figs 1 to 8 have been obtained by numerically solving the ordinary differential equations provided in A1 to A4 via the odeint function from the integrate module of the Scientific Python (SciPy) package with a constant stepsize dt = 0.01 h. The Matplotlib library has been used to generate figures. Codes are available upon request.

### 4.2. Calculation of Hill coefficients and delays

An ultrasensitive response is often sigmoidal and the curve can be well approximated by the Hill equation. The effective Hill equation is defined for the temporal curves in Figs 2, 3 and 6 (see Equation 1). The effective Hill coefficient n is related to the effective time 90% (ET90) and 10% (ET10) ratio by Equation 2. Here K is the effective time when 50% (or ET50) of total phosphorylation is achieved.

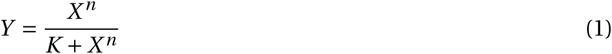

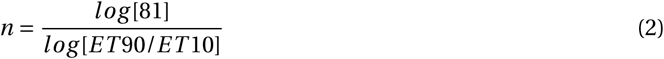

Delays are calculated as the time when 50% of the maximal value is achieved for each phosphoprotein Fk as given in Equation 3.

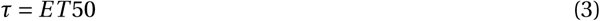

## Author Contributions

conceptualization by M.B. and H.H.; methodology by A.U. and H.H.; software by A.U.; validation by A.U. and H.H.; formal analysis by A.U.; investigation, A.U. and H.H.; resources by H.H.; data curation by M.B., D.M. and A.D.; writing—original draft preparation by A.U. and H.H.; writing—review and editing, A.U., H.H., D.M., A.D. and M.B.; visualization by A.U. and H.H.; supervision by M.B. and H.H.; project administration by M.B. and H.H.; funding acquisition by M.B. and H.H.

## Acknowledgments

The authors are thankful to Christoph Schmal and Bharath Ananthsubramaniam for fruitful discussions.

## Appendix (Supporting Information)

### Appendix A.1 Scheme of the linear model, equations and parameters

We modelled the linear phosphorylations using five-variable ordinary differential equations (ODEs). Mass-action kinetics of processive phosphorylations provide long delays shown in Fig 2 with the parameters k=50 and *k*_*d*_ =0.1.

**Figure A1.**
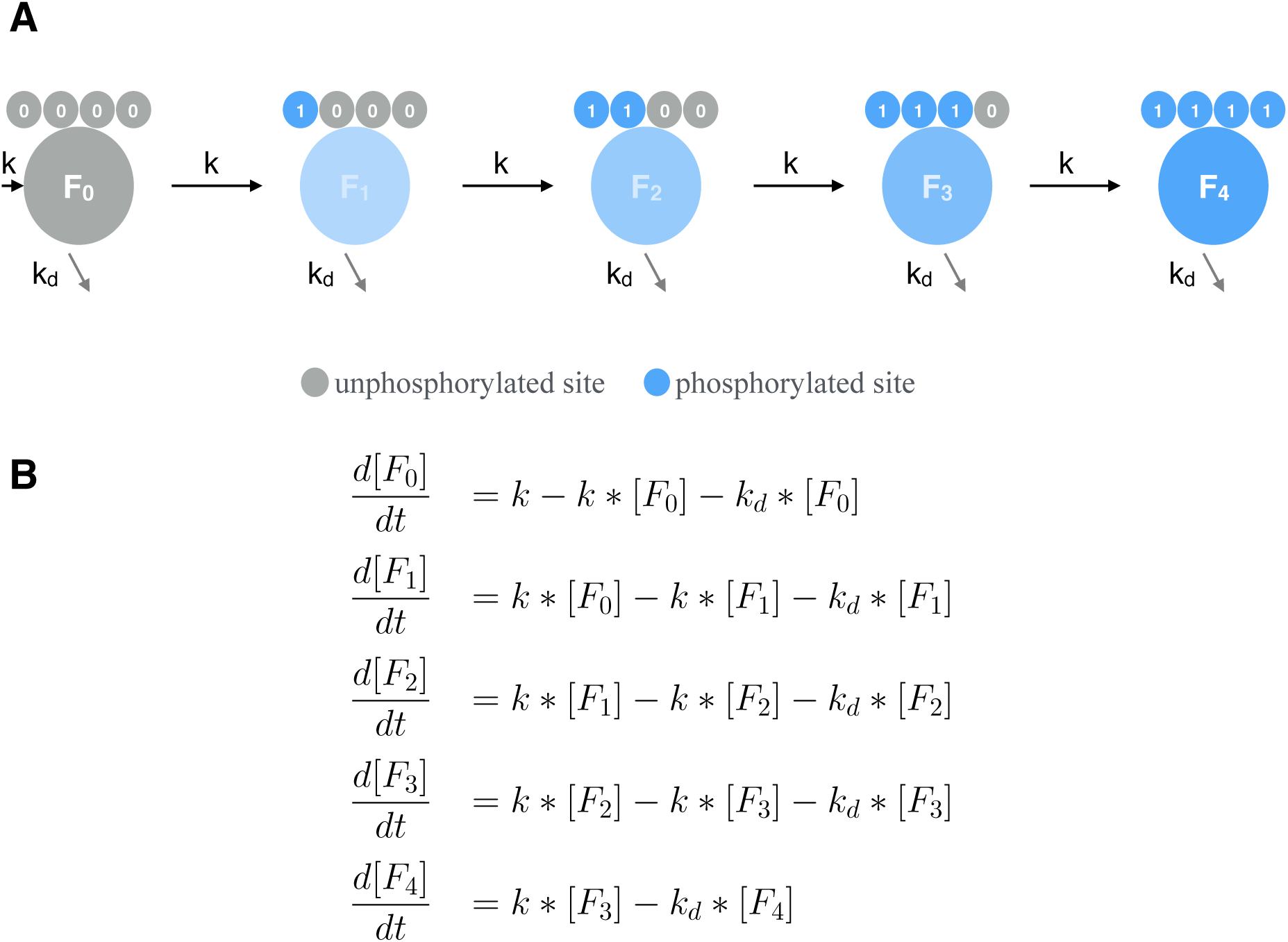
**(A) Linear model: Schematic diagram of phosphorylations with turnover of protein F:** F is phosphorylated in a sequential processive manner. **(B) Model of up to 4 phosphorylations with 5 variables**.

### Appendix A.2 Scheme of the nonlinear model, equations and parameters

We modelled the nonlinear phosphorylations using a ten-variable ODE system. Distributive kinetics of phosphorylations enhances ultrasensitivity shown in Fig 3. Parameters in Figs 3 and 5: *k*_1_=50, *k*_2_=50, *k*_*d*_ =0.1, *P*_*p*_ =500 and *P*_*c*_ =1. Fig 4: *P*_*c*_ =5.

**Figure A2.**
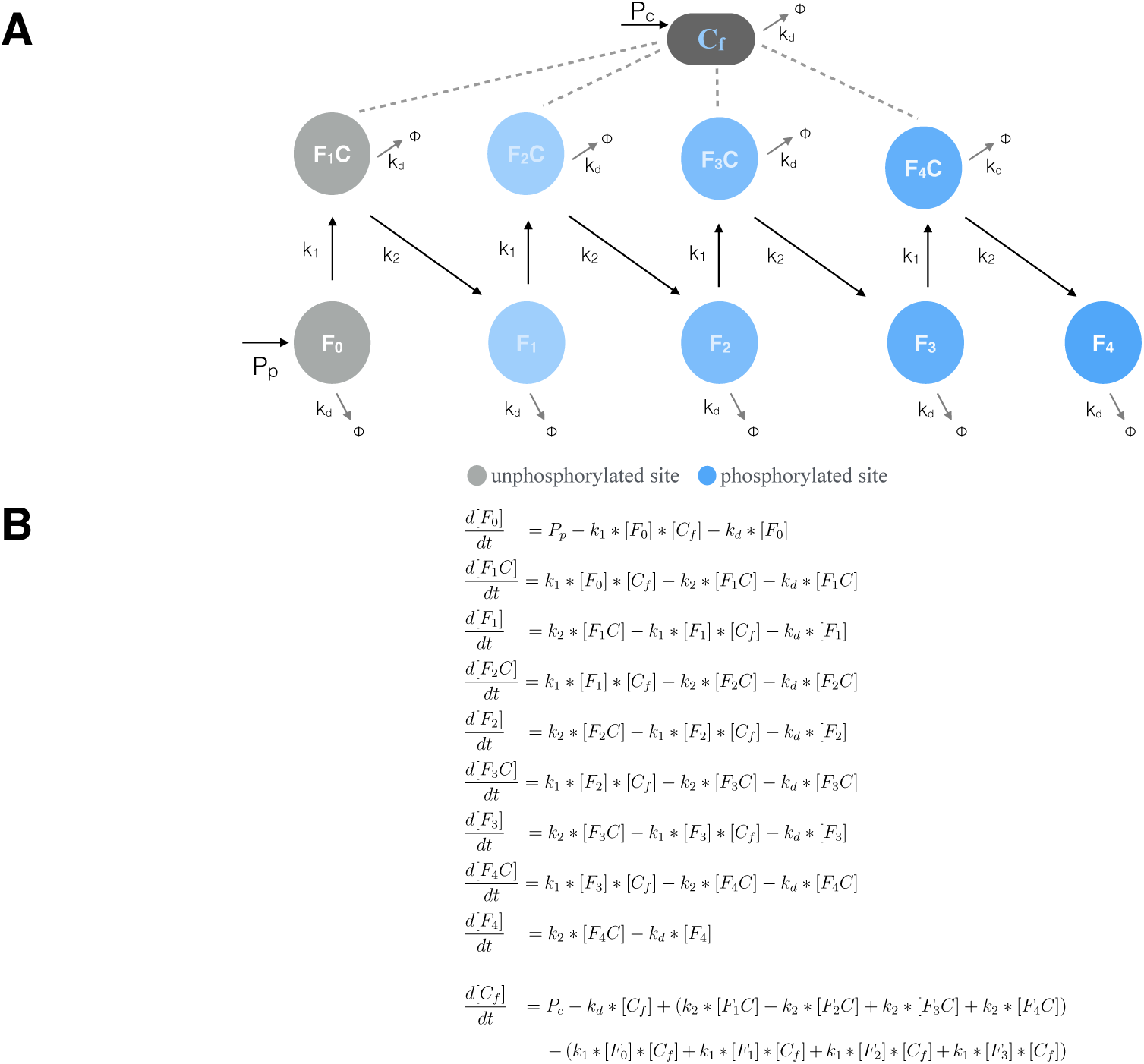
**(A) Nonlinear model: Schematic diagram of protein phosphorylations with turnover of F and C:** F is phosphorylated by C in a sequential distributive manner. **(B) Model of up to 4 phosphorylations of F with 10 variables**.

### Appendix A.3 Noninear random model: scheme, equations and parameters

We modelled the nonlinear random phosphorylations using a ten-variable ODE system. Distributive kinetics and prefactors associated to random phosphorylations together provide large amplitudes of phosphorylations shown in Fig 6. Parameters in Fig 6: *k*_1_=50, *k*_2_=50, *k*_*d*_ =0.1, *P*_*p*_ =500 and *P*_*c*_ =1.

**Figure A3.**
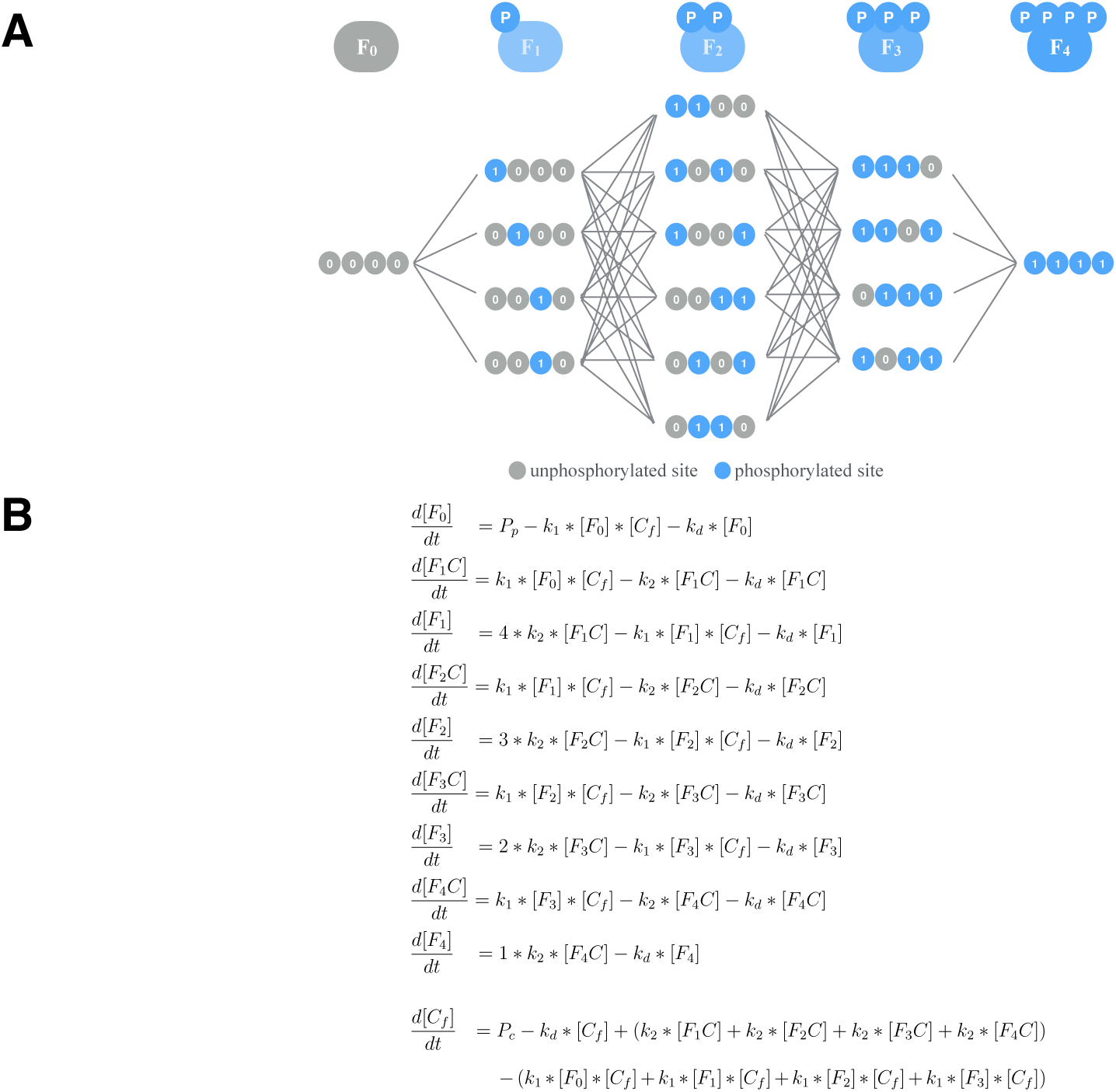
**(A) Nonlinear random model: Schematic diagram of phosphorylations with turnover of protein F and kinase C:** F is phosphorylated by C in a distributive random manner. **(B) Model of up to n=4 phosphorylations**.

### Appendix A.4 Large scale nonlinear random model: scheme, equations and parameters

We modelled the large scale nonlinear random phosphorylations using a 2n+2 (n=100) variable ODE system. Distributive kinetics, prefactors associated to random phosphorylations and longer chain of phosphorylations are shown in Fig 7. Amplitudes, delays and Hill coefficients for 100 phosphorylations shown in Fig 8 are also derived from model simulations. Parameters in Fig 7: *k*_1_=50000, *k*_2_=50000, *k*_*d*_ =1.5, *P*_*p*_ =0.01 and *P*_*c*_ =10. Default (blue) parameters in Fig 8: *k*_1_=50000, *k*_2_=50000, *k*_*d*_ =5, *P*_*p*_ =1 and *P*_*c*_ =10. Slow degradation (red): *k*_*d*_ =0.15 and high kinase/protein ratio (green): *P*_*c*_ =100.

**Figure A4.**
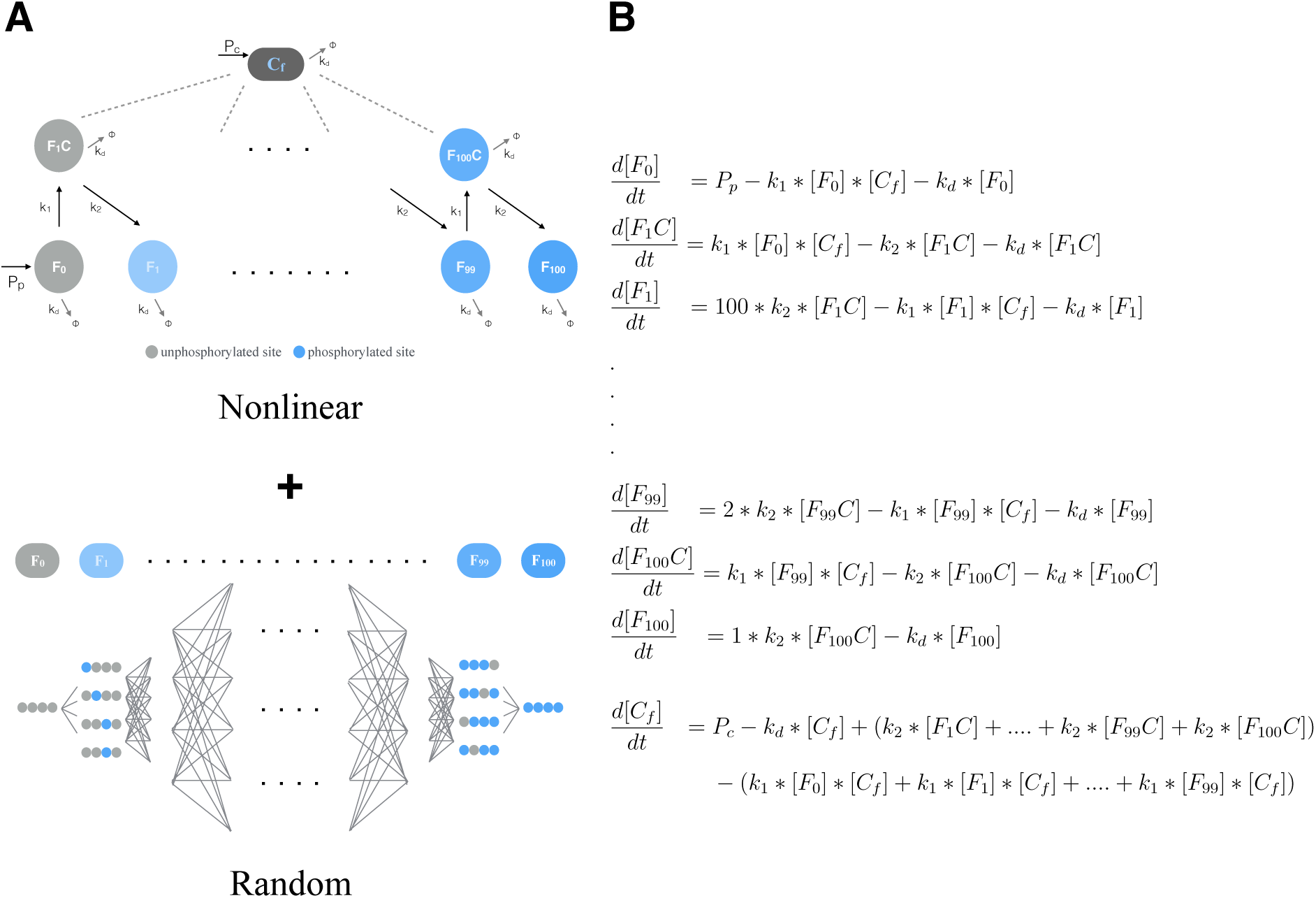
**(A) Large scale nonlinear random model: Schematic diagram of phosphorylations with turnover of protein F and kinase C:** F is phosphorylated by C in a distributive random manner. **(B) Model of up to n=100 phosphorylations with 2n+2 variables**.

### Appendix A.5 Hill coefficients and delays

Hill coefficients and delays are calculated using eqns 1, 2 and 3 for the curves shown in Figs 2, 3 and 6.

**Figure A5.**
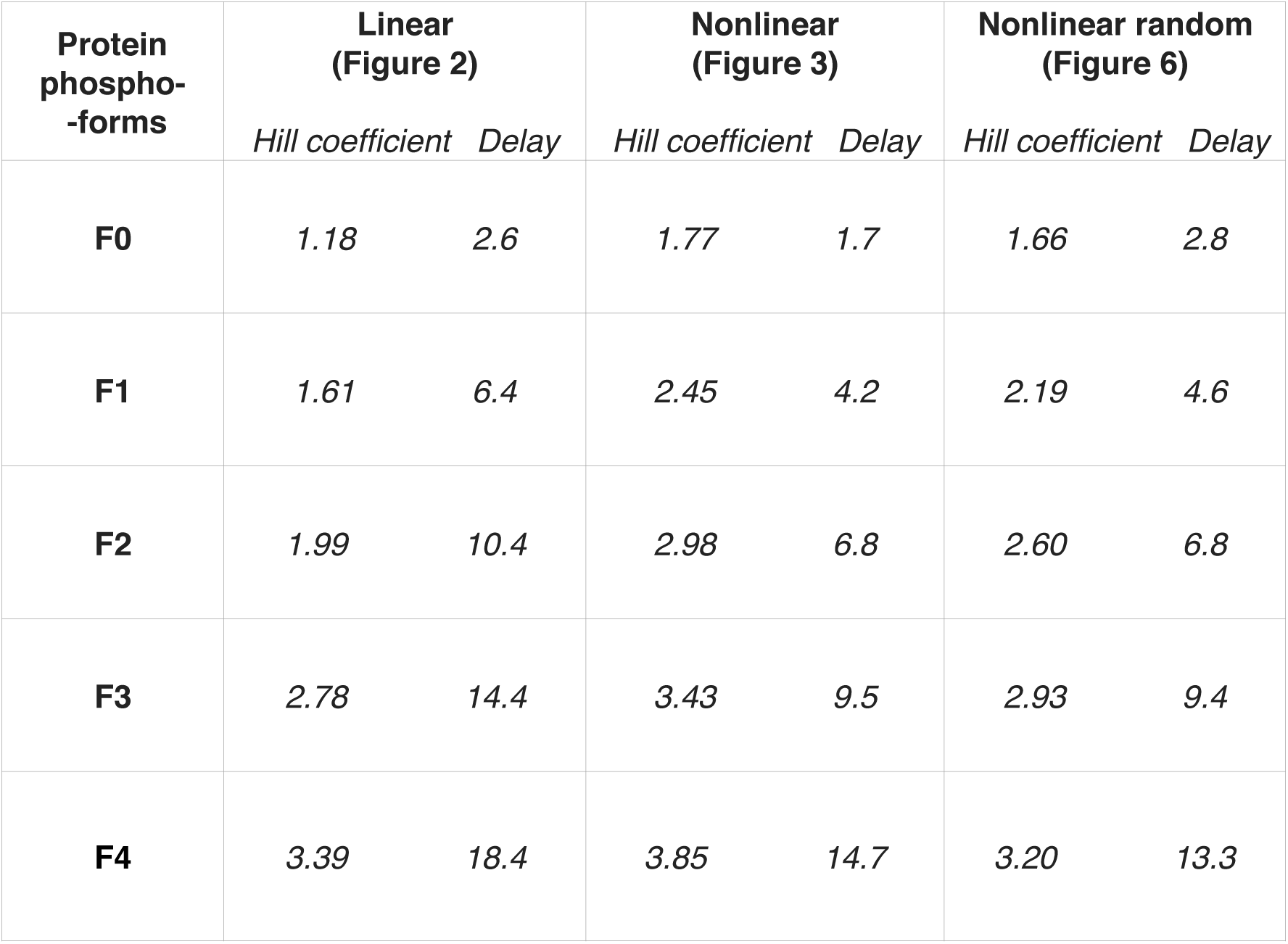
Hill coefficients and delays: A list of Hill coefficients and time delays of the phosphorylated Fk across three models. See Figs 2, 3 and 6.

### Appendix A.6 Enlarged time series of nonlinear phosphorylations

**Figure A6.**
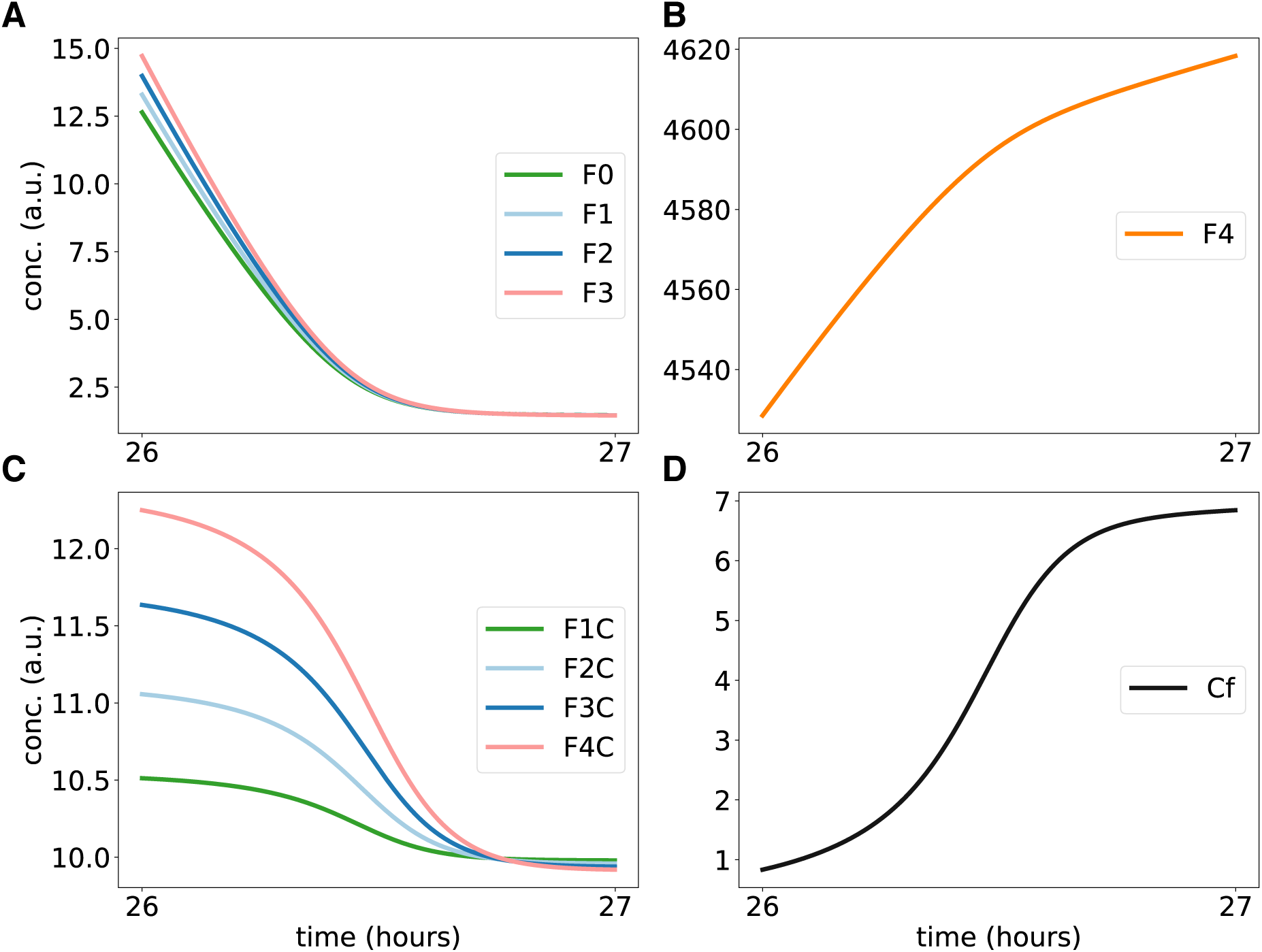
Magnified simulations of nonlinear phosphorylations from Fig 4: Graphs show a one hour time window (26 to 27 hours) and confirm that apparent kinks in Fig 4 are rather smooth curves. The asymptotic equilibrium indicates that most of the F protein is sequestered within complexes.

### Appendix A.7 Effects of model extensions

**Figure A7.**
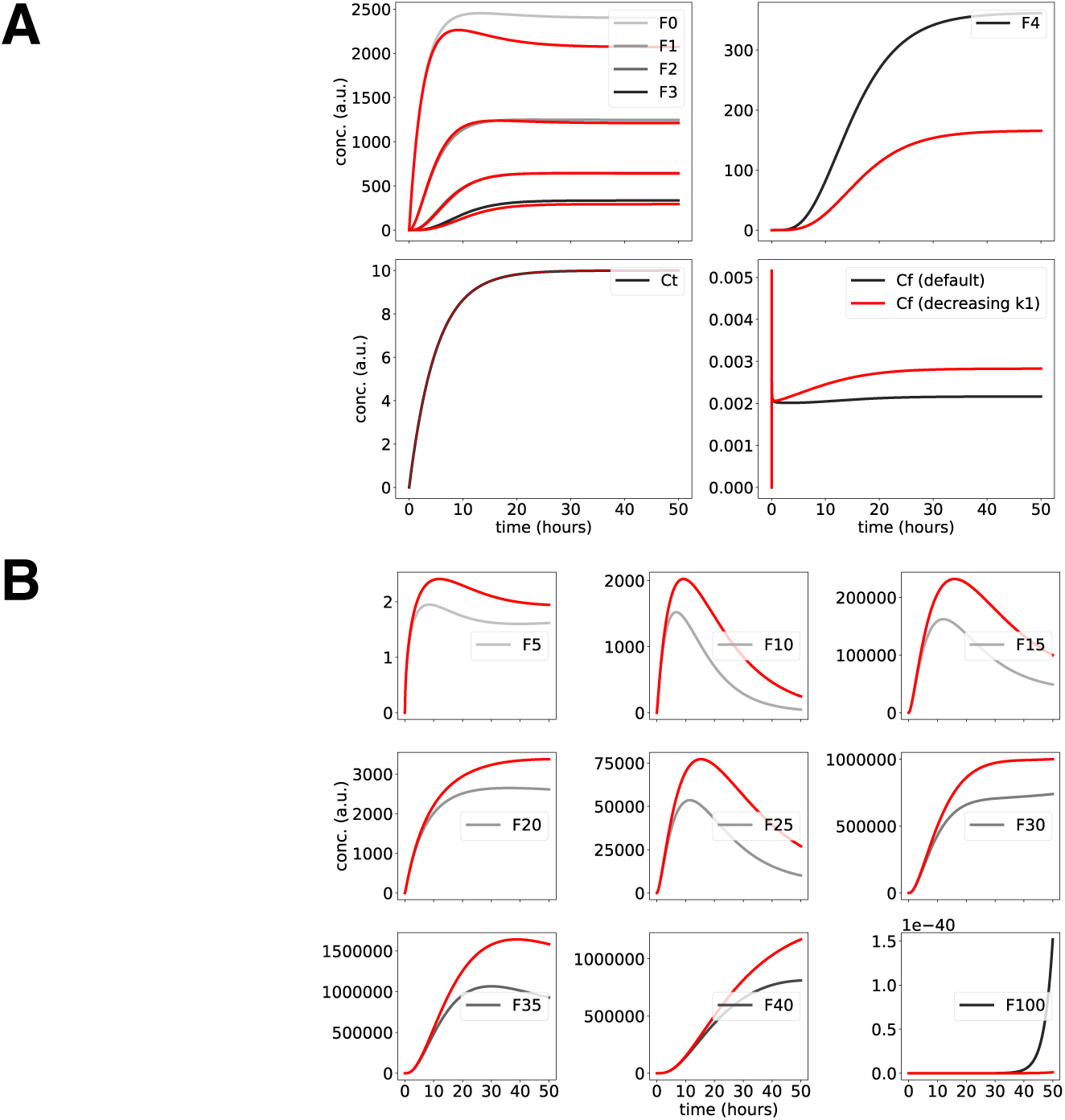
**(A) Simulating decreasing phosphorylation rates:** We compare constant phosphorylation rates *k*_1_ (see Fig 3 and A2) with decreasing rates over increasing phosphorylation: 50, 40, 30, 20. The modifications have only minor effects on amplitudes, delays, and waveforms. **(B) Simulating stabilizing and destabilizing phosphorylations:** Compared to constant degradation rates *k*_*d*_ =1.5 (grey curves) we take into account the stabilizing effects of initial phosphorylation and the destabilization of later phosphorylations. We decrease for k=0, 1,…, 50 *k*_*d*_ by a factor 0.95 and increase *k*_*d*_ for k=51, 52,…, 100 by 0.95^−1^. This modification leads to even higher amplitudes of intermediate phosphorylation levels (red curves).

